# Integrated Transcriptomic and Functional Analysis Reveals Tissue-Specific Molecular Pathology in Adolescent Idiopathic Scoliosis

**DOI:** 10.64898/2026.05.27.727643

**Authors:** Darius Ramkhalawan, Paola Parrales, Justin Koesterich, Gloria Montoya-Vazquez, Carlos Cuna, Anat Kreimer, Jessica McQuerry, Stephanie Ihnow, Nadja Makki

## Abstract

Adolescent idiopathic scoliosis (AIS), the spontaneous development of a lateral spine curvature during puberty, is the most common pediatric spine disorder, affecting ∼3% of children worldwide. As the underlying etiology remains unclear, AIS is treated purely symptomatically, initially by bracing and ultimately by highly invasive, costly surgeries. Genome-wide association studies (GWAS) have identified numerous risk loci in non-coding genomic regions, making it difficult to link them to a biological function. To address this, we performed a multi-tissue investigation to connect genetic risk to tissue-specific molecular pathology.

We conducted RNA sequencing on the primary tissues implicated in AIS, paraspinal muscle and spinal cartilage, from patients and unaffected controls. In paraspinal muscle, we identified differentially expressed genes (DEGs) enriched for pathways related to muscle structure, myogenesis, and metabolism. Key upregulated genes include the transcription factor *EGR1* and structural components such as *MYH1*. In spinal cartilage, we found enrichment of genes related to TGFβ and FoxO signaling, as well as metabolic pathways. Notably, genes crucial for chondrocyte differentiation (e.g. *SOX5* and *SOX6*) were significantly downregulated. We then examined genes at known GWAS loci and found that several risk-associated genes were differentially expressed in one or both tissues. To investigate the function of non-coding variants at these loci, we identified and validated several enhancer elements harboring AIS risk SNPs at the *BCL2, ADGRG6, BNC2, and FTO* loci. We reveal distinct pathological signatures in muscle and cartilage and lay the foundation for connecting non-coding genetic risk to the dysregulation of key developmental and structural pathways.

## Introduction

Adolescent idiopathic scoliosis (AIS [MIM: 181800]) is a lateral curvature of the spine that spontaneously develops and progresses during the adolescent growth spurt^1^. Current treatments for AIS are limited to restrictive bracing in moderate cases, or corrective surgery by way of implantation of metal instruments adjacent to the patient’s spine in severe cases^2^. AIS affects approximately 3% of children worldwide, however its etiology remains unclear, and it is currently not possible to identify predisposed patients before symptoms are clinically evident. Due to the poor understanding of the mechanisms underlying curve onset and progression, AIS must be treated symptomatically, as there are no preventative measures or pharmacological treatment options.

A genetic component of AIS has been well-established through reported family histories and twin studies^3^. Population-specific genome-wide association studies (GWAS) and meta-analyses have identified a number of genetic susceptibility loci^4–18^. In addition, animal studies that knocked out specific genes or gene regulatory elements have resulted in scoliosis-like phenotypes^8,19–23^, revealing that several tissues are involved in AIS pathogenesis, including paraspinal muscle and spinal cartilage. Further, histological and electrophysiological studies of paraspinal muscle biopsies reported asymmetry in muscle fiber size, fiber type, and electrical activity between the concave and convex side of the curvature^24–27^. Cartilage-specific conditional gene knockout models have resulted in scoliosis-like phenotypes^19^. Cumulatively, these studies reveal a complex genetic architecture of AIS where multiple genes acting in different tissues, possibly in an asymmetric fashion, contribute to the phenotype. However, previous efforts to characterize the AIS transcriptome have been limited by a lack of control samples. Thus, a major goal for AIS research is to identify the gene regulatory networks that are affected in the different tissues and ultimately understand how genetic variants affect these networks. Since most GWAS variants reside in non-coding regions, one goal is to identify and validate the activity of specific gene regulatory elements, such as enhancers, that harbor these AIS-associated variants. Such information will be key for predicting risk, prognosis, treatment response, and clinical outcomes.

To better understand the genetic changes that occur in AIS tissues, we performed transcriptional profiling and differential expression analysis of two key tissues involved in AIS pathogenesis: paraspinal muscle and spinal cartilage. Our analysis identified previously unreported genes related to muscle differentiation, function, and structure, as well as genes related to chondrocyte differentiation and extracellular matrix, as being differentially expressed between AIS patients and unaffected controls. In addition to profiling gene expression, we identified several enhancer elements that harbor AIS-associated variants and validated their allele-specific regulatory activity in relevant cell types. Our integrated analysis of differential gene expression and enhancer activity in AIS tissues reveals distinct molecular signatures of disease progression, highlighting key pathways and regulatory elements that can be further investigated as potential biomarkers and therapeutic targets. Ultimately, this study provides a foundational genomic resource that ties genetic risk to tissue-specific dysfunction, opening avenues for mechanistic research into AIS onset and progression.

## Subjects, material, and methods

### Tissue collection

All methods in this study were performed in accordance with the guidelines and regulations outlined in a protocol approved by the University of Florida Institutional Review Board (Study ID: IRB202003156). 16 patients between the ages of 12 and 20 undergoing AIS corrective surgery were assessed by an orthopaedic surgeon and formal written consent was obtained from the parent/guardian to donate paraspinal muscle and facet joint cartilage. Equivalent muscle samples were collected from 5 control individuals undergoing spinal surgery for unrelated reasons. Detailed information on subjects included in this study can be found in Table S1. Specimens were flash-frozen in liquid nitrogen prior to RNA isolation.

### RNA extraction

Frozen tissues were pulverized in liquid nitrogen using a mortar and pestle, followed by homogenization in TRIzol (ThermoFisher Scientific, cat no. 15596026) using a bead beater (TissueLyser LT, Qiagen, cat no. 69980) at 50 Hz for 1 min with 1/8” stainless steel beads (MP Biomedicals, cat no. 1169250-CF), followed by RNA isolation, per the TRIzol manufacturer’s protocol.

### RNA sequencing and analysis

Messenger RNA was purified from total RNA using poly-T oligo-attached magnetic beads. After fragmentation, the first strand cDNA was synthesized using random hexamer primers followed by the second strand cDNA synthesis. Paired-end RNA sequencing was carried out on an Illumina NovaSeq6000.

Control cartilage RNA-seq data was accessed from Chen et al. and originate from one male and two female vertebral fracture patients between the ages of 18 and 32 undergoing internal fixation of the lumbar spine (detailed sample information is available in Table S1). In brief, Chen and colleagues separated facet joint cartilage from subchondral bone, snap froze tissue in liquid nitrogen, pulverized the tissue and extracted RNA with TRIzol reagent (Invitrogen), followed by purification with the RNeasy Mini kit (Qiagen). RNA sequencing was carried out on an Illumina HiSeq X10 instrument.

Briefly, we utilized the nf-core’s RNA-seq pipeline with the selection to pseudo-align the reads to genes with the program Salmon. We next loaded the Salmon count matrices into a DESeq dataframe using the Tximport R package^51^ and generated the comparison model as such. The DESeq model accounted for age, sex, and disease status (AIS or control)^52^. Finally, we outputted the log fold change and adjusted p-values for each gene and identified differential genes with log2FC ≥ 1 or ≤ -1 and an adjusted p-value threshold ≤ 0.05.

### Data and code availability

Raw sequencing data for all AIS and control muscle is available from the Gene Expression Omnibus under accession GSE311180. AIS spinal cartilage data is available from the Sequence Read Archive under accession PRJNA625649. Healthy cartilage control data is available at the European Nucleotide Archive under study accession PRJNA474389.

### Gene enrichment analyses

GO analysis was performed with on DEGs using the clusterProfiler R library. Raw p-values were adjusted using Benjamini-Hochberg correction, then redundant terms were collapsed with the simplify function using the Wang Semantic Similarity method. Significant terms were selected using p-value cutoff of 0.05 and q-value cutoff of 0.1. KEGG analysis was performed using ShinyGO 0.85^53–55^ which interfaces with the Ensembl and STRING-db databases to extract enrichment information from gene sets and removed redundant terms that shared 95% of their genes and 50% of the words in their names.

### Luciferase reporter assays

Candidate enhancers were amplified from human genomic DNA (Promega, cat no. G304A) by PCR and cloned into pGL4.23[luc2/mP] (Promega, cat no. E8411) using the NEBuilder HiFi DNA assembly kit (NEB, cat no. E2621S), then transformed into DH5α competent cells (NEB, cat no. C2987H) and plated on carbenicillin selection plates overnight at 37°C. Successful insertion of the CRS was verified by colony PCR, plasmids were isolated by miniprep (Macherey-Nagel, cat no. 740490.50), and sequenced. Cells were plated in 24-well plates to yield 80% confluency the next day. For C2C12 myoblasts, cells were transfected using EndoFectin Max (GeneCopoeia, cat no. EF013) 24 hours after seeding. The next day, the culture media was replaced with differentiation media (DMEM supplemented with 2% heat-inactivated horse serum) and cells were allowed to differentiate for 72 hours. For TC28a2 chondrocytes, cells were transfected with 900 ng of the plasmid and 100 ng of pGL4.74[hR/luc] (Promega, cat no. E6921) using X-tremeGENE HP DNA Transfection Reagent (Roche, cat no. 6366244001) according to the manufacturer’s protocol. After 48 hours post-transfection, luciferase activity was assayed using the Dual Luciferase Reporter Assay System (Promega, cat no. E1980) according to the manufacturer’s protocol on a Glomax Multi Detection System plate reader (Promega, cat no. 9301-010).

### Prediction of disrupted transcription factor binding motifs

Transcription factor binding site disruptions were predicted using the motifbreakR program^56^. MotifbreakR utilizes a position weight matrix from the ENCODE database of transcription factor binding sites to predict if a variant causes a neutral, weak, or strong disruption to a predicted binding site. We only considered “strong” disruptions with p ≤ 1×10^-4^ in this study.

### Statistics

Statistically significant DEGs were identified using the DESeq2 package, which normalizes read counts and calculates an adjusted p-value using the Wald test with Benjamini-Hochberg correction. For luciferase assays, each construct was compared to empty vector negative controls (pGL4.23) with at least 3 biological replicates and statistical significance was determined by one-way ANOVA with Dunnett’s test to account for multiple comparisons. Statistical difference between alleles was determined with at least 6 biological replicates and significance was calculated by multiple unpaired t-tests with Benjamini-Krieger-Yekutieli correction for multiple comparisons. Differences between undifferentiated and differentiated cell states were determined using single unpaired Student’s t-tests.

## Results

### Differential expression analysis in paraspinal muscle identifies genes with high relevance to AIS pathogenesis

As paraspinal muscle and vertebral cartilage are the two major tissues implicated in AIS pathogenesis^19,22,24,25,27–33^, we carried out differential RNA-seq analysis on both tissues as a crucial step for connecting GWAS-identified risk loci to the functional changes in gene expression that drive the deformity. RNA was isolated from the paraspinal muscle and facet joint cartilage from AIS patients undergoing spinal fusion surgery (Table S1). Samples from AIS patients formed a distinct cluster, separate from controls as shown by principal component analysis (PCA; Figure 1A). Samples were primarily separated by condition (i.e. AIS or control), however within groups, samples from the same sex tended to cluster together, reflecting the sexually dimorphic nature of AIS.

**Figure 1.**
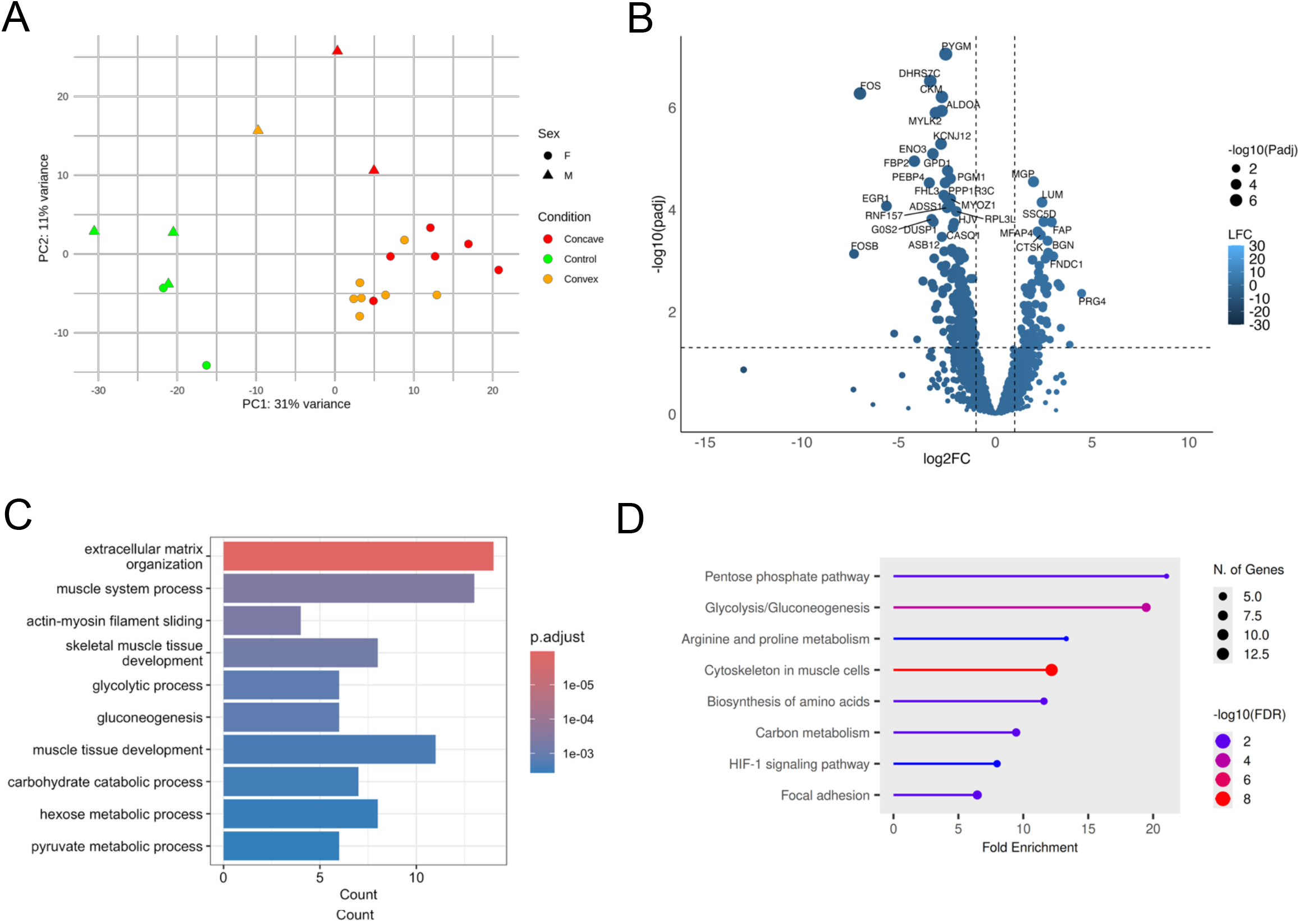
Differential expression of AIS concave paraspinal muscle and functional enrichment of DEGs. A) Principal component analysis of paraspinal muscle samples dissected from AIS patients (n=10) and controls (n=5). B) Volcano plot of DEGs between AIS concave and control muscle. Larger dot size indicates greater statistical significance, lighter blue indicates more positive log2 Fold Change and darker blue indicates more negative log2 Fold Change. Horizontal dashed line indicates padj = 0.05, vertical dashed lines indicate log2FC = -1 and log2FC = 1. C) Gene ontology (GO) annotation of DEGs between concave and control muscle samples. D) KEGG pathway enrichment analysis^55^ of differentially expressed genes in concave versus control muscle samples displaying the top 10 enriched KEGG pathways.

We first examined expression in paraspinal muscle by comparing the concave (n=8) and convex (n=8) sides of the curve to identify asymmetrically expressed genes. We identified 101 differentially expressed genes (DEGs), which were primarily enriched for muscle cytoskeletal components (Figure S1, Table S2, and Table S3).

We next compared expression of each side of the curve to unaffected adult paraspinal muscle as controls (n=5). We identified 300 and 567 DEGs (log2FC ≥ 1 or ≤ -1, padj ≤ 0.05) when comparing muscle from the concave side (Figure 1B, Table S4) and convex (Figure 2A, Table S5) sides of the curve, respectively, to control tissues. To identify pathways with strong biological effect, we limited gene enrichment analyses to include only high-confidence DEGs with log2FC ≥ 2 or ≤ -2 and padj ≤ 0.05. This resulted in 108 DEGs on the concave side, and 224 DEGs on the convex side.

**Figure 2.**
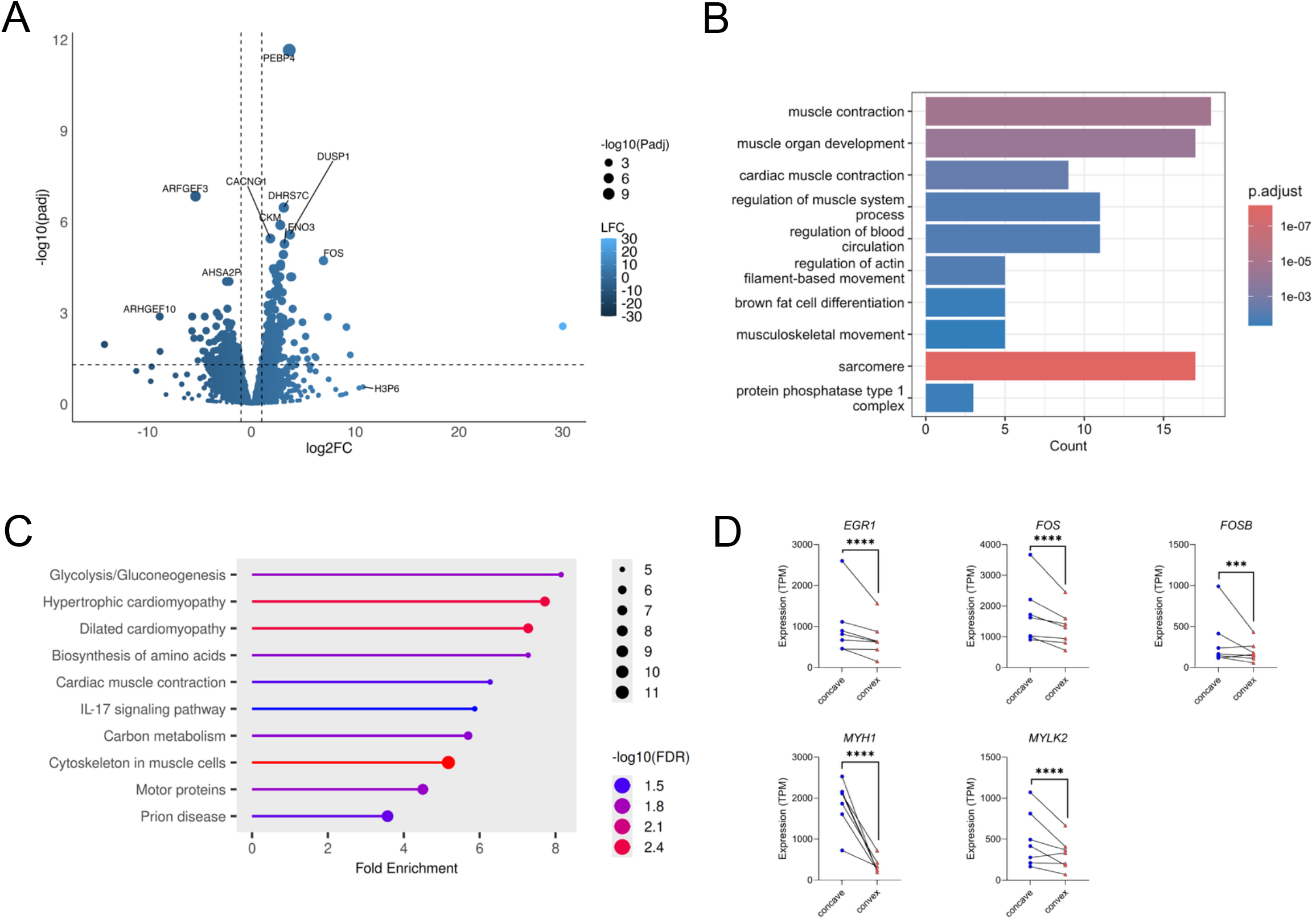
Differential expression of AIS convex paraspinal muscle and functional enrichment of DEGs. A) Volcano plot of DEGs between AIS convex and control muscle. Larger dot size indicates greater statistical significance, lighter blue indicates more positive log2 Fold Change and darker blue indicates more negative log2 Fold Change. Horizontal dashed line indicates padj = 0.05, vertical dashed lines indicate log2FC = -1 and log2FC = 1. Gene ontology (GO) annotation of DEGs between concave and control muscle samples. KEGG pathway enrichment analysis^55^ of differentially expressed genes in concave versus control muscle samples displaying the top 10 enriched KEGG pathways. D) Selected genes that are asymmetrically expressed within individual patients.

Gene ontology (GO) annotation of DEGs between concave and control muscle revealed enrichment of genes related to extracellular matrix organization and muscle tissue development (Figure 1C, Table S6). DEGs between the convex and control muscle, were enriched for terms such as “muscle contraction,” “sarcomere,” and “musculoskeletal movement” (Figure 2B, Table S7). KEGG pathway enrichment analysis of DEGs in concave versus control muscle also identified several metabolic pathways (pentose phosphate, glycolysis/gluconeogenesis, biosynthesis of amino acids, carbon metabolism); cytoskeleton in muscle cells, and focal adhesion as the top 10 enriched pathways (Figure 1D, Table S8). DEGs on the convex side were largely enriched in pathways related to muscle function, such as “cytoskeleton in muscle cells” and “motor proteins” (Figure 2C, Table S9)

Among the top DEGs between AIS and control muscle is early growth response 1 (*EGR1* [MIM: 128990]), which is upregulated in AIS muscle compared to controls, and asymmetrically expressed, being more highly expressed on the concave side of patients (Figure 2D). *EGR1* encodes a transcription factor that was shown to transactivate an enhancer of *UNCX*, which contains the AIS risk SNP rs78148157^21^, and has been reported to regulate satellite cell differentiation during skeletal muscle development^34,35^. Also among the most highly upregulated and asymmetrically expressed genes are *FOS* [MIM: 164810] and *FOSB* [MIM: 164772], which encode transcription factors known to promote myogenesis^36,37^. Several genes encoding structural myofibril components, which are essential for muscle fiber formation and contraction, were found to be differentially expressed between AIS and control muscle and asymmetrically expressed, including myosin 1 (*MYH1* [MIM: 160730]), myosin light chain kinase 2 (*MYLK2* [MIM: 606566]), and skeletal alpha actin (*ACTA1* [MIM: 102610]).

### Expression of AIS risk loci in paraspinal muscle

GWAS have identified dozens of AIS risk loci, however, their relevance to AIS and the tissues they affect are still largely unknown ^4,6–9,11–13,18,19,22,36^. Our RNA-seq analysis has allowed us to measure the expression of genes at the associated loci and identify those that are differentially expressed in paraspinal muscle (Figure 3A). We included genes proximal to all currently reported GWAS loci (Wise et al. (2020)^38^), as this provides a pre-defined list of GWAS loci, several of which have been replicated in multiple studies. In total we found 19/29 genes expressed in AIS muscle on both the concave and convex sides of the curve (defined as average TPM ≥ 1), while one gene (*ABO* [MIM: 110300]) was expressed only on the convex side.

**Figure 3.**
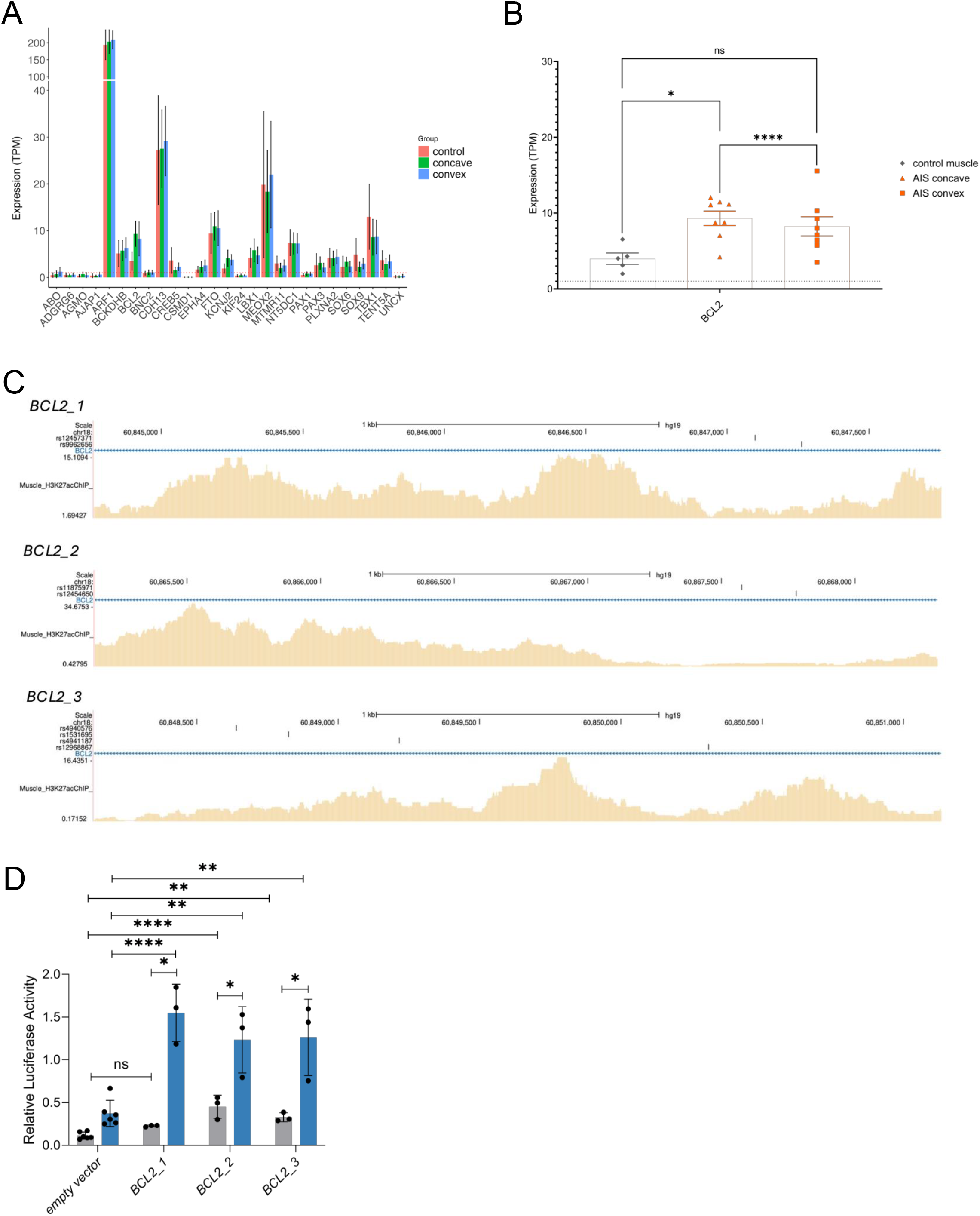
Expression of AIS-associated genes and myocyte enhancers at the *BCL2* locus. A) Expression of AIS-associated genes in paraspinal muscle of AIS patients and controls. Dotted line indicates TPM = 1. B) Expression of differentially expressed AIS-associated genes in paraspinal muscle of AIS patients and controls. Statistical significance was determined using a Wald test on normalized DESeq counts followed by Benjamini-Hochberg correction. Error bars indicate mean ± SD. Dotted line indicates TPM = 1. C) Enhancers at the *BCL2* locus that are active in undifferentiated myoblasts (grey bars) and differentiated myotubes (blue bars). Statistically significant enhancer activity relative to empty vector negative controls was calculated using one-way ANOVA with Dunnett’s test, comparing either each candidate enhancer to empty vector negative controls, or activity in the undifferentiated to the differentiated cell state. Error bars indicate mean ± SD. NS, not significant, *, p < 0.05, **, p < 0.01, ****, p < 0.0001 E) Functional enhancers at the *BCL2* locus overlap with H3K27ac enrichment in AIS patient muscle (yellow) and contain AIS risk SNPs.

Of the examined genes at the AIS risk loci, only *BCL2* [MIM: 151430] was differentially expressed between the concave and convex side and relative to control samples, where it was upregulated in AIS concave versus control (log2FC = 1.4; Figure 3B). *ABO*, which was expressed only on the convex side of the curve (average TPM = 1.22), but not the concave side (average TPM = 0.68), was not considered differentially expressed, most likely due to the read counts being too low and therefore being filtered out by DESeq2.

### Identification of functional enhancers at the *BCL2* locus

The lead variant at the *BCL2* locus, rs4940576, as well as all variants in linkage disequilibrium (r^2^ ≥ 0.8) are located in intron 2 of the *BCL2* gene (MANE Select NM_000633.3). Comparing the location of these variants with H3K27ac ChIP data derived from AIS patient muscle generated by Makki et al.^31^ showed significant levels of H3K27ac enrichment (a marker of active enhancers) in intron 2 of *BCL2*. Given their location in a non-coding sequence and significant levels of H3K27ac enrichment at this locus, we hypothesized that associated variants at the *BCL2* locus may be affecting enhancer elements. To test this, we cloned three individual regions surrounding rs4940576 (Figure 3C) into a luciferase reporter vector upstream of a minimal promoter and transfected the constructs into C2C12 myoblasts. We identified significant levels of enhancer activity relative to empty vector negative controls in undifferentiated myoblasts, which increased further post-differentiation (Figure 3D). These enhancers are distributed throughout the linkage block (Figure 3C), suggesting the presence of several adjacent enhancers centered around rs4940576. Several SNPs in this region are predicted to disrupt transcription factor binding motifs which have relevance to muscle biology (Figure S2). For example, rs12457371, located in the *BCL2_1* enhancer is predicted to alter OSR1 binding (Figure S2A), which mediates activation of fibroadipogenic progenitor cells during muscle injury^39,40^.

### Differentially expressed genes in AIS cartilage are enriched for key signaling pathways that regulate chondrocyte homeostasis

To identify differentially expressed genes in vertebral cartilage, the other major tissue implicated in AIS pathogenesis, we performed expression profiling of cartilage from the facet joints of AIS patients^31^. Due to the limited amounts of cartilage that can be harvested from patients, tissue from the concave and convex sides of the patient’s curvature were pooled together prior to RNA isolation. As we could not acquire equivalent tissue from unaffected controls, we identified a previously published dataset of facet joint cartilage from healthy patients^41^ and performed differential expression analysis. AIS cartilage clustered separately from controls by PCA (Figure 4A). We identified 3,360 upregulated (log2FC ≥ 1, padj ≤ 0.05) genes and 3,069 downregulated (log2FC ≤ -1, padj ≤ 0.05) genes in AIS cartilage (Figure 4B, Table S10). For downstream analyses, we used high confidence DEGs with log2FC ≥ 2 or ≤ -2 and padj ≤ 0.05, resulting in 2,396 upregulated genes and 2,097 downregulated genes. GO terms were related to ATP synthesis and muscle cytoskeletal components, which could be indicative of chondrocyte dedifferentiation^42^ or mechanical stress^43^ (Figure 4C, Table S11). KEGG pathway enrichment showed overrepresentation of pathways related to stress and immune responses (e.g. Coronavirus disease), TGFβ signaling, FoxO signaling, focal adhesion, and ECM-receptor interaction (Figure 4D, Table S12).

**Figure 4.**
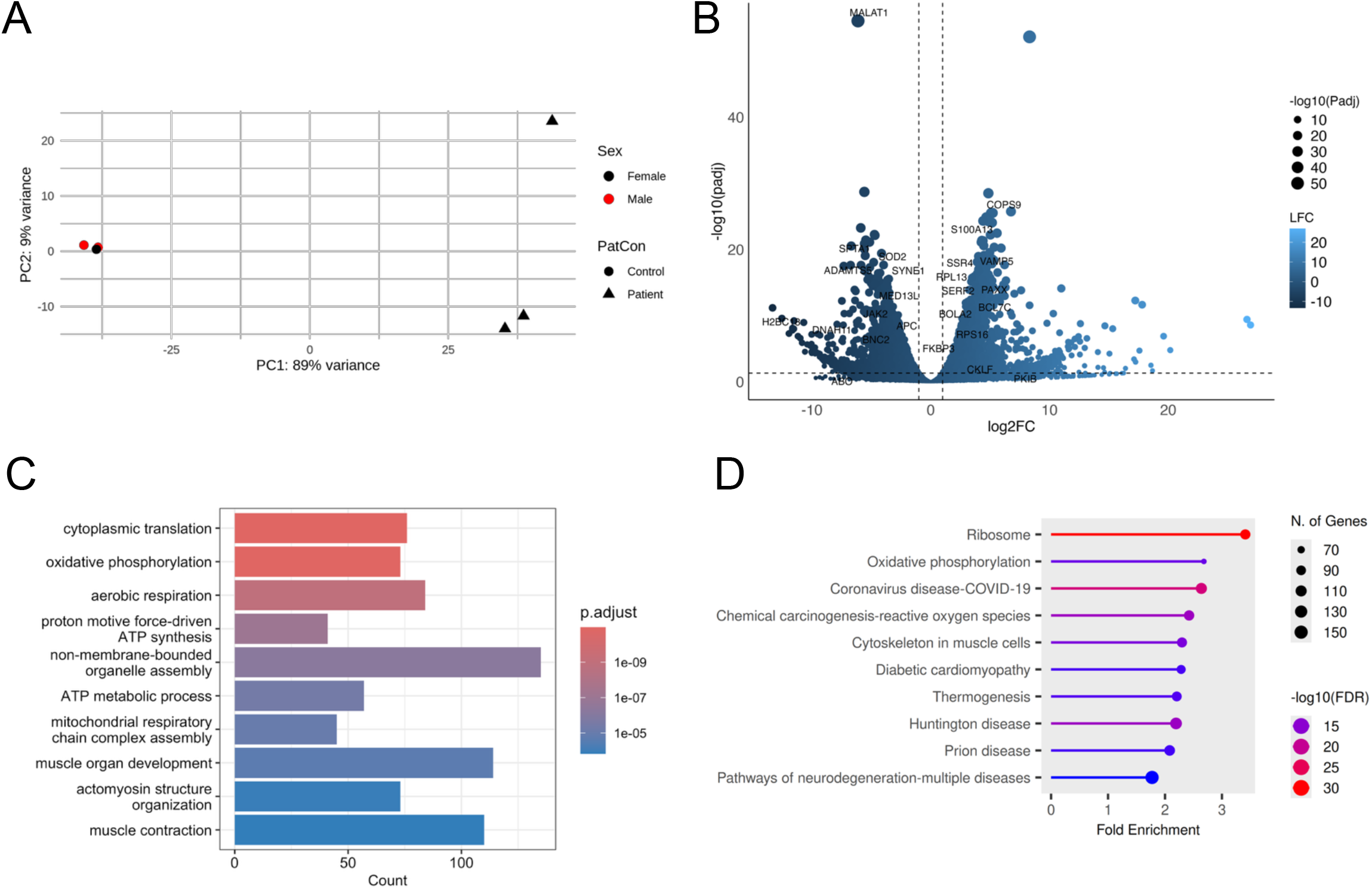
Differential expression of AIS spinal cartilage and functional enrichment of DEGs. Principal component analysis of spinal cartilage samples dissected from AIS patients (n=3) and controls (n=3). B) Volcano plot of DEGs between AIS and control cartilage. Larger dot size indicates greater statistical significance, lighter blue indicates more positive log2 Fold Change and darker blue indicates more negative log2 Fold Change. Horizontal dashed line indicates padj = 0.05, vertical dashed lines indicate log2FC = -1 and log2FC = 1. C) GO annotation of DEGs between AIS and control cartilage samples. D) KEGG pathway enrichment analysis of differentially expressed genes in AIS versus control cartilage samples displaying the top 20 enriched KEGG pathways.

### Genes associated with AIS through GWAS are differentially expressed in spinal cartilage

We identified 15/29 genes at AIS associated loci to be expressed in AIS spinal cartilage (average TPM ≥ 1; Figure 5A). Differential expression analysis identified 7/29 as being differentially expressed between AIS and healthy spinal cartilage samples (Figure 5B). Several of these differentially expressed loci have high relevance to cartilage biology in their role of regulating chondrocyte differentiation and ECM production, such as the adhesion molecule *ADGRG6* [MIM: 612243] and the transcription factor *SOX6* [MIM: 607257]. Other differentially expressed loci include *ARF1* [MIM: 103180], *ABO, BNC2* [MIM: 608669], *FTO* [MIM: 610966], *and PAX1* [MIM: 167411].

**Figure 5.**
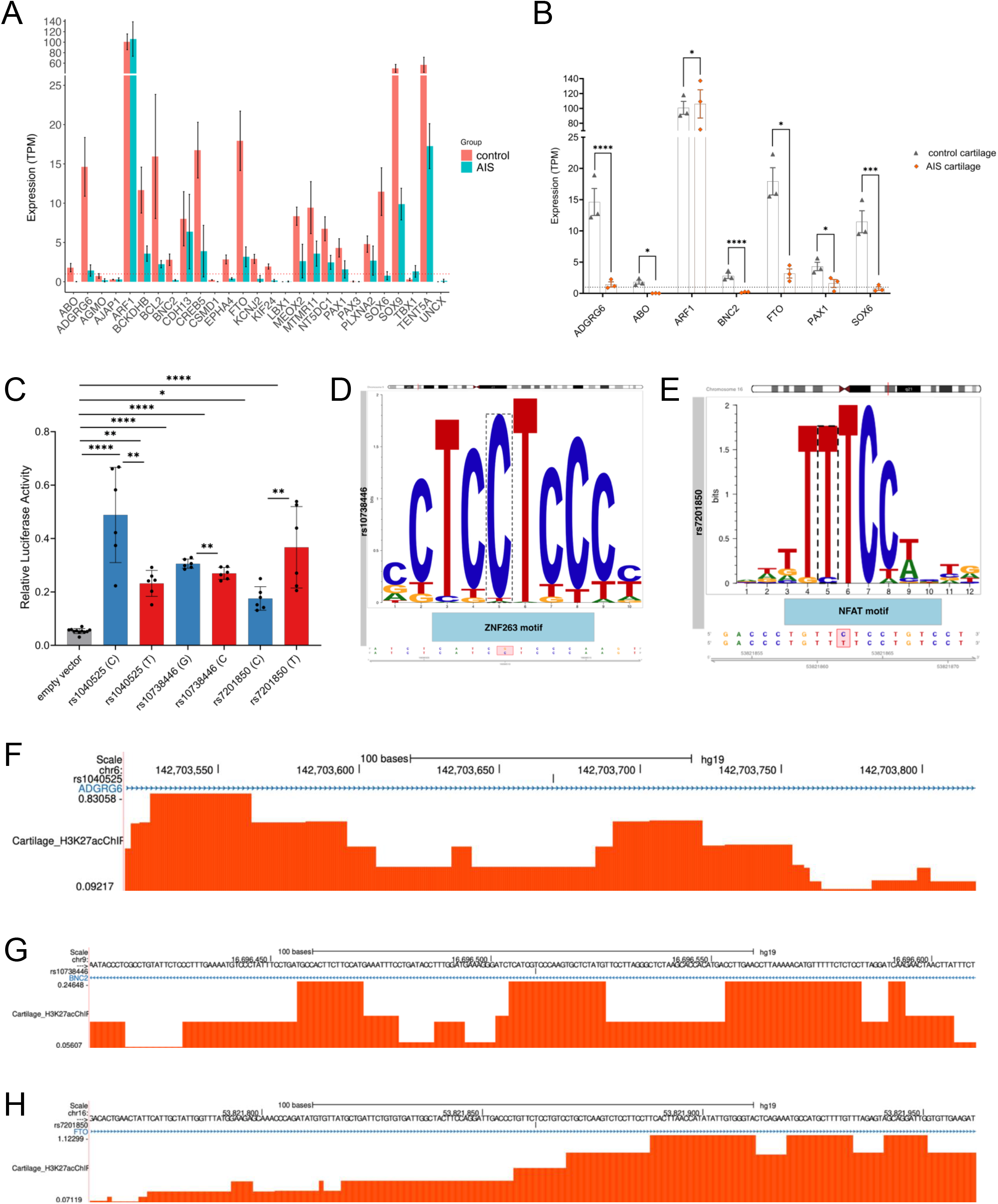
Expression of AIS-associated genes and identification of chondrocyte enhancers at the *ADGRG6, BNC2*, and *FTO* loci. A) Expression of AIS-associated genes in spinal cartilage of AIS patients (n=3) and controls (n=3). Dotted line indicates TPM = 1. B) Expression of differentially expressed AIS-associated genes in spinal cartilage of AIS patients and controls. Statistical significance was determined using a Wald test on normalized DESeq counts followed by Benjamini-Hochberg correction. Error bars indicate mean ± SD. Dotted line indicates TPM = 1. C) Enhancers at the *ADGRG6, BNC2, and FTO* loci that are active in chondrocytes. Statistical significance comparing candidate enhancer to empty vector negative controls was calculated using one-way ANOVA with Dunnett’s test; statistical differences between reference and alternate alleles of the same SNP were calculated using multiple unpaired Student’s t-test followed by Benjamini-Krieger-Yekutieli correction for multiple comparisons. Error bars indicate mean ± SD. D) Predicted transcription factor binding motif disruption caused by rs10738446. E) Predicted transcription factor binding motif disruption caused by rs7201850. F) rs1040525 enhancer locus showing AIS facet joint cartilage H3K27ac enrichment (red). G) rs10738446 enhancer locus showing AIS facet joint cartilage H3K27ac enrichment (red). H) rs7201850 enhancer locus showing AIS facet joint cartilage H3K27ac enrichment (red).

### Identification of functional SNPs at the *ADGRG6, BNC2, and FTO* loci

We identified chondrocyte enhancers at the *ADGRG6, BNC2*, and *FTO* loci that harbor AIS associated variants and showed allele-specific differences in enhancer activity relative to empty vector negative controls (Figure 5C). The enhancer at the *ADGRG6* locus is located in intron 5 of the *ADGRG6* gene (MANE Select NM_198569.3) and contains SNP rs1040525 (Figure 5F). The enhancer at the *BNC2* locus is located in intron 3 of *BNC2* (MANE Select NM_017637.6) and contains the risk SNP rs10738446, which is predicted to alter ZNF263 binding, where the risk allele (G) completely abolishes the TFBS (Figure 5D, G). Finally, we identified an enhancer at the *FTO* locus which harbors the SNP rs7201850, which is located in intron 1 of the *FTO* gene (MANE Select NM_001080432.3), and is predicted to affect NFAT binding, where the T allele is highly preferred in the binding sequence (Figure 5E, H). ENCODE eQTL data indicates that rs1040525 targets *ADGRG6* in several biosamples; rs10738446 targets *BNC2-AS1*, the antisense RNA of *BNC2*; and rs7201850 targets *FTO* (Table S12).

## Discussion

AIS is a complex, polygenic disorder, thus, identification of the involved genes and their interaction in common networks and pathways is key to the understanding of AIS etiology. In addition to the complex genetic nature of this disorder, several tissues have been implicated in AIS pathogenesis, making it important to study disease-associated genes and gene regulatory networks in a tissue-specific manner.

To identify tissue-specific expression profiles, we first carried out RNA-seq analysis on two of the tissues most relevant to AIS pathology. GO analysis of differentially expressed genes in AIS vs control muscle revealed enrichment of genes related to muscle structure and function. Finally, KEGG pathway enrichment analysis shows that DEGs in muscle are related to metabolic pathways, such as the pentose phosphate pathway and gluconeogenesis. The etiology of AIS is multifactorial and not well understood. Various mechanisms have been investigated as the potential root of its etiology, such as altered energy metabolism^44^ and calcium levels^32^. Our analysis suggests that other metabolic dysfunctions may also be occurring in patients.

One of the top DEGs between AIS and control muscle is *EGR1*, which is upregulated in AIS muscle on the convex side and downregulated on the concave side. *EGR1* encodes a transcription factor that was shown to transactivate an enhancer of *UNCX*, which contains the AIS risk SNP rs78148157^21^. EGR1 shows a higher binding affinity for the risk allele of rs78148157 resulting in upregulated *UNCX* expression in neuronal cells. It will be interesting to investigate if patients that carry the AIS risk SNP rs78148157 at the *UNCX* locus also show an upregulated expression of *EGR1*, which would suggest either a positive feedback loop, or independent mechanisms that lead to an upregulation of *UNCX* in neurons. *EGR1*, as well as several of the other most significant DEGs between AIS patients and controls (e.g. *FOS, FOSB*), plays a key role during myogenesis. In addition to genes regulating myogenesis, we identified several genes encoding structural myofibril components, which are essential for muscle fiber formation and contraction, to be highly differentially expressed between AIS and control muscle, including genes encoding myosin heavy and light chains, myosin binding proteins, and troponins. Asymmetries in paraspinal muscle fiber type and function on the convex and concave side of the curve have long been reported in the literature as a possible mechanism underlying AIS pathogenesis^25,30,45,46^.

When comparing AIS and control cartilage, DEGs were enriched for pathways related to TGFβ and FoxO signaling. In the context of cartilage, they are responsible for different aspects of chondrocyte differentiation and homeostasis: TGFβ signaling represses chondrocyte differentiation and maintenance and FOXO promotes chondrocyte maturation^47,48^. Genes related to focal adhesion and ECM-receptor interaction were also enriched in our data set, indicating impaired ECM composition and integrity may play a role in the onset and progression of AIS. Genes related to chondrocyte differentiation, such as *SOX5* [MIM: 604975] and *SOX6*, were downregulated in patients compared to controls. GO term analysis showed enrichment of genes related to ATP synthesis, such as “oxidative phosphorylation”. Dedifferentiated chondrocytes may have a preference for oxidative phosphorylation and may undergo cytoskeletal and ECM reorganization^42^.

In addition to identifying genes involved in AIS pathogenesis, a major goal of our study was to investigate the expression profiles of genes previously associated with AIS through GWAS. In paraspinal muscle, we identify *BCL2* as being upregulated on the concave side of the curve. *BCL2* was reported to be expressed during early myogenesis and promote clonal expansion of myoblasts^49^. Its asymmetrical expression may have implications for AIS curve formation and/or progression. As all AIS GWAS SNPs at the *BCL2* locus are located in intron 2, we set out to identify enhancer elements overlapping associated SNPs. We identified 3 enhancer elements that demonstrated significant luciferase activity both pre- and post-differentiation into myotubes, where regulatory activity increased significantly post-differentiation. These elements are distributed throughout the linkage block, suggesting the presence of several adjacent enhancers centered around rs4940576. Our transcription factor binding site (TFBS) analysis of this region predicted that several SNPs could disrupt motifs relevant to muscle biology. For example, the rs12457371 SNP, located within the *BCL2_1* enhancer element, is predicted to alter an OSR1 binding site. Given OSR1’s role in mediating the activation of fibroadipogenic progenitor cells during muscle injury, this finding is particularly noteworthy. The precise functional consequences of this predicted interaction, and its potential disruption by local SNPs, will need to be investigated further.

In cartilage we identified several genes at AIS loci to be differentially expressed between AIS and control, namely *ABO, ADGRG6, ARF1, BNC2, FTO, PAX1*, and *SOX6*. Several other genes were trending toward differential expression, however potentially due to limited statistical power, differences were not significant. As all GWAS variants at the above loci are in non-coding regions, we carried out enhancer analyses at three of the loci: *ADGRG6 BNC2*, and *FTO*. Our functional analyses provide a potential mechanism for these non-coding variants. We demonstrated that AIS-associated SNPs at the *ADGRG6, BNC2*, and *FTO* loci reside within active chondrocyte enhancers and drive allele-specific activity (Figure 5C). The *ADGRG6* variant (rs1040525) appears to directly target *ADGRG6* expression. This is particularly compelling given that mouse models lacking *Adgrg6* in osteochondral progenitors develop a scoliosis-like phenotype^19,29^, supporting its role in AIS pathogenesis. Similarly, the *BNC2* risk variant (rs10738446) likely alters gene expression by affecting ZNF263 binding, with eQTL data pointing to the antisense RNA *BNC2-AS1* as the regulatory target. At the *FTO* locus, we identified the functional SNP rs7201850, which targets the *FTO* gene in gastrocnemius muscle, however its target in chondrocytes has not yet been identified. FTO is an RNA demethylase that removes m6A modifications from a variety of RNA molecules. Recent studies in osteoarthritis models have found there is an overabundance of m6A in osteoarthritic human cartilage, and that overexpressing *FTO* can alleviate symptoms and protect cartilage from degradation^50,51^ Taken together, our findings suggest these non-coding AIS risk variants may contribute to pathogenesis by functionally impacting allele-specific expression of key target genes in chondrocytes.

### Limitations

Our study provides insights that will inform future mechanistic studies to dissect the intricate interplay of different genetic loci and different spinal tissues in driving susceptibility and progression of this complex disease. In our study we used healthy adult paraspinal muscle and spinal cartilage as controls due to the limited availability of age-matched tissues, as these tissues are rarely removed from healthy adolescent children. Future differential expression analyses with age-matched tissues will be necessary to draw more definitive conclusions regarding tissue-specific pathology.

As cartilage is a major tissue implicated in AIS etiology, further examination of spinal cartilage is of particular interest for further study, though several major challenges exist. Whereas paraspinal muscle is abundantly available, and previous studies have identified structural differences between muscle from the concave and convex sides of the scoliotic curve, similar analyses for spinal cartilage are lacking. Spinal cartilage from individuals with AIS tends to be arthritic, and therefore in smaller quantities, due to the asymmetric mechanical stress on the spine. This poses a technical challenge in acquiring sufficient tissue for DE analysis of concave and convex cartilage. In this study, we pooled cartilage from the concave and convex sides of the curve to identify genetic pathways that may be driving curve onset. Further studies are required to identify specific differences between the cartilage of concave and convex sides of the curve. We were unable to acquire healthy spinal cartilage as controls. To compensate, our cartilage DE analysis drew on samples from a previously published source of healthy adult spinal cartilage, which has the potential to introduce batch effects. While batch was taken into account in the DE analysis, our limited sample size limits statistical power, and therefore larger sample sizes will be required to gain further biological insights.

### Future Directions

To determine how these genes might affect AIS pathogenesis, it is crucial to determine which tissues express them, whether they show differential expression, and which other genes act in common pathways or networks. Differential expression of myogenesis genes (e.g. *FOS, BCL2*) suggests that myogenesis is somehow impaired or otherwise affected in AIS, possibly in an asymmetric manner. Further experiments at the protein level and in independent cohorts comparing the differentiation potential and apoptotic activity of muscle from the concave and convex sides of the curve and unaffected muscle could yield insights into how asymmetric cell survival and myogenic ability contribute to muscle remodeling, and ultimately progression of AIS. It should also be noted that muscle tissue is also comprised of several different cell types, including satellite cells and fibroadipogenic progenitor cells (FAPs). Analysis of these cell types, in addition to myoblasts and myocytes, could have implications for understanding the origin of AIS, making single-cell sequencing an important next step in understanding AIS etiology.

We also find differential expression of genes related to chondrocyte differentiation. It is interesting to speculate that asymmetric chondrocyte differentiation triggers onset of the curve, which is then exacerbated as the child grows. Animal models would be especially useful to test the developmental consequences of aberrant chondrocyte differentiation and hypertrophy. Given the shared developmental origin of muscle and cartilage in mesenchymal stem cells (MSCs), further study of MSCs derived from AIS patients is warranted.

Finally, we identify several active enhancers that harbor AIS risk variants and demonstrate allele-specific differential enhancer activity. Additional validation of enhancer activity in primary cells derived from AIS tissues will help establish the biological relevance of these enhancers. Further functional analysis, for example with CRISPR activation or inhibition, is also needed to link these enhancers to their downstream target genes and to identify their role in disease etiology. Several risk variants were also predicted to affect transcription factor binding; validating these effects through ChIP or mobility shift assays would help establish regulatory pathways affected in AIS. Our findings therefore present a preliminary, though important, link between genetic risk variants and changes in gene expression in AIS tissues.

## Supplemental Information Description

Supplemental information contains two figures and 16 tables.

## Author Contributions

N.M. and D.R. conceived the experiments. S.I. and J.M. recruited and evaluated patients and collected human tissue samples. D.R., G.M-V., P.P., and C.C. performed experiments. D.R., J.K., A.K., and N.M. analyzed the data. N.M. and D.R. wrote the manuscript.

## Acknowledgements

We would like to thank Dr. Robert Decker, Dr. Nigel Price, Johanna Carmona, Camesha Tate, and Dr. Jennifer Shumway for their assistance in recruiting patients and collecting tissues for this study. We are grateful to Dr. Rhonda Bacher for input on statistical analysis. This work was funded by the Scoliosis Research Society (SRS) and the University of Florida Research Opportunity Seed Fund awarded to N.M., S.I., and J.M.

## Figure Legends

**Figure S1**. Volcano plot of differentially expressed genes between the concave (n=8) and convex (n=8) sides of the scoliotic curve.

**Figure S2**. A) Predicted transcription factor binding motif disruption caused by rs12457371. B) Predicted transcription factor binding motif disruption caused by rs9962656. C) Predicted transcription factor binding motif disruption caused by rs4941187. D) Predicted transcription factor binding motif disruption caused by rs12968887.

## Table Legends

**Table S1**. Tissue donor demographic information.

**Table S2**. Concave muscle vs convex muscle DESeq2 results.

**Table S3**. Concave muscle vs convex muscle GO term enrichment results.

**Table S4**. Concave muscle vs control muscle DESeq2 results.

**Table S5**. Convex muscle vs control muscle DESeq2 results.

**Table S6**. Concave muscle vs control muscle GO term enrichment results.

**Table S7**. Convex muscle vs control muscle GO term enrichment results.

**Table S8**. Concave muscle vs control muscle KEGG enrichment results.

**Table S9**. Convex muscle vs control muscle KEGG enrichment results.

**Table S10**. AIS cartilage vs control cartilage DESeq2 results.

**Table S11**. AIS cartilage vs control cartilage GO term enrichment results.

**Table S12**. AIS cartilage vs control cartilage KEGG enrichment results.

**Table S13**. eQTL data from RegulomeDB.

**Table S14**. Gene TPM values.

**Table S15**. Primer sequences used to generate CRSs.

**Table S16**. Raw firefly and renilla luciferase values.

