## Supplemental Figures and Tables for "Integrated Transcriptomic and Functional Analysis Reveals Tissue-Specific Molecular Pathology in Adolescent Idiopathic Scoliosis": Supplemental Figures and Table S15.pdf

Figure S1

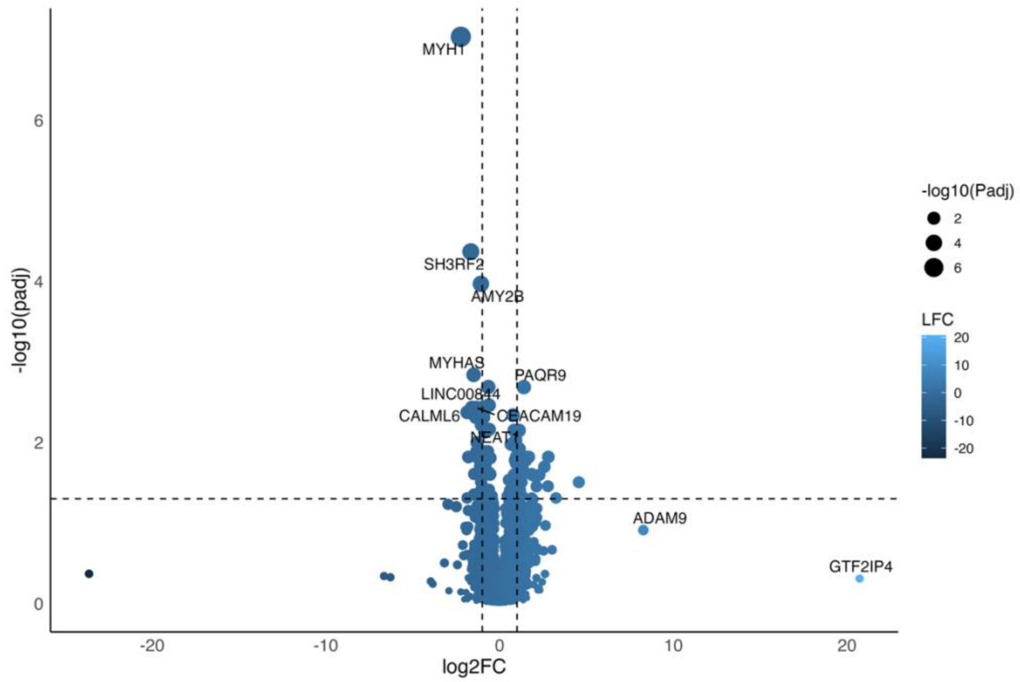

MYH1
SH3RF2
AMY2B
MYHAS
PAQR9
LINC00844
CALML6
CEACAM19
NEAT1
ADAM9
GTF2IP4
-log10(padj)
log2FC
-log10(Padj)
LFC

### Figure S2

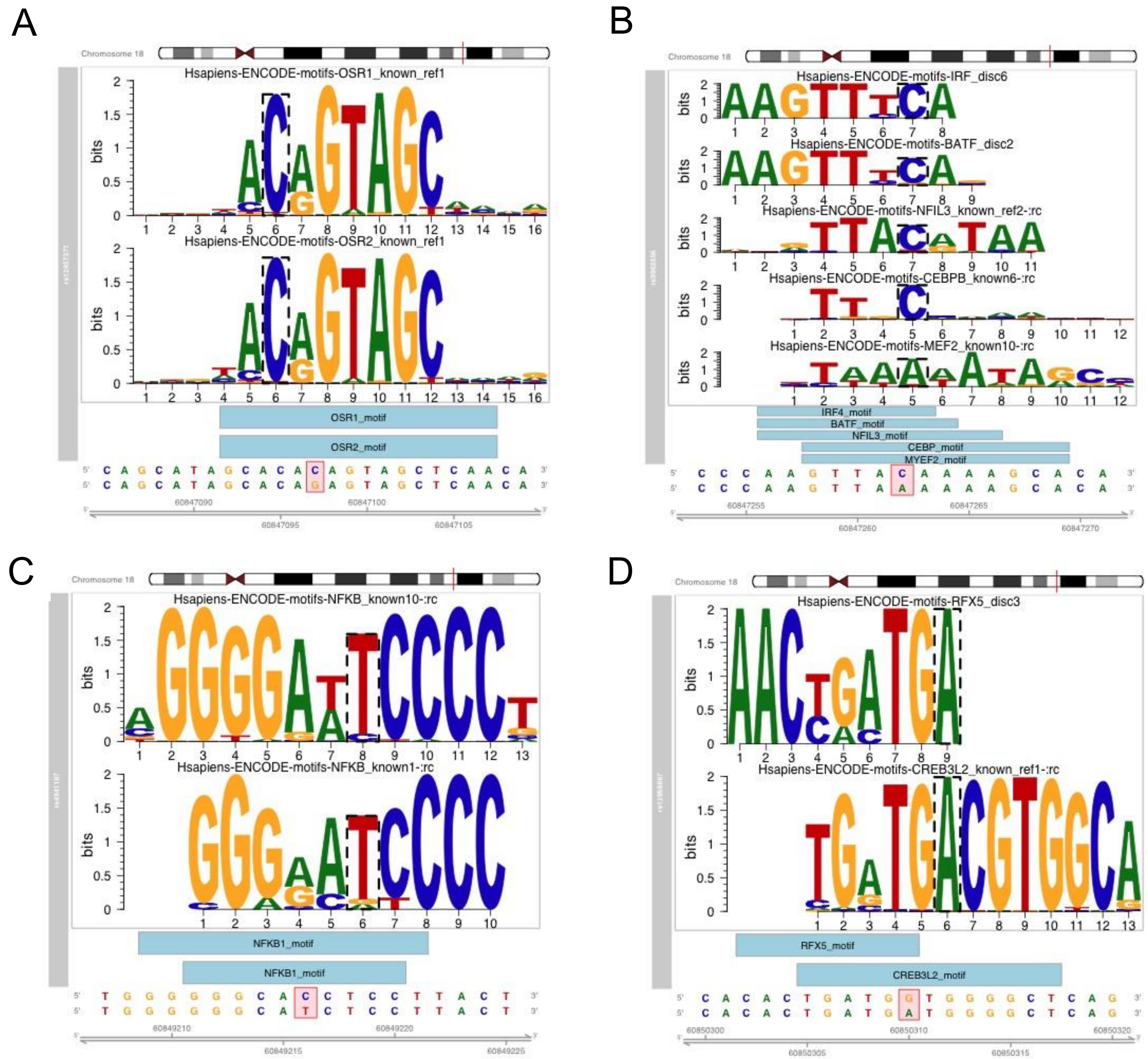
A
Chromosome 18
Hsapiens-ENCODE-motifs-OSR1_known_ref1
Hsapiens-ENCODE-motifs-OSR2_known_ref1
bits
OSR1_motif
OSR2_motif
C A G C A T A G C A C A C A G T A G C T C A A C A
C A G C A T A G C A C A G A G T A G C T C A A C A
60847090
60847100
60847095
60847105
B
Chromosome 18
Hsapiens-ENCODE-motifs-IRF_disc6
Hsapiens-ENCODE-motifs-BATF_disc2
Hsapiens-ENCODE-motifs-NFIL3_known_ref2-:rc
Hsapiens-ENCODE-motifs-CEBPB_known6-:rc
Hsapiens-ENCODE-motifs-MEF2_known10-:rc
bits
IRF4_motif
BATF_motif
NFIL3_motif
CEBP_motif
MYEF2_motif
C C C A A G T T A C A A A A G C A C A
C C C A A G T T A A A A A A G C A C A
60847255
60847265
60847260
60847270
C
Chromosome 18
Hsapiens-ENCODE-motifs-NFKB_known10-:rc
Hsapiens-ENCODE-motifs-NFKB_known1-:rc
bits
NFKB1_motif
NFKB1_motif
T G G G G G G C A C C T C C T T A C T
T G G G G G G C A T C T C C T T A C T
60849210
60849220
60849215
60849225
D
Chromosome 18
Hsapiens-ENCODE-motifs-RFX5_disc3
Hsapiens-ENCODE-motifs-CREB3L2_known_ref1-:rc
bits
RFX5_motif
CREB3L2_motif
C A C A C T G A T G G T G G G G C T C A G
C A C A C T G A T G A T G G G G C T C A G
60850300
60850310
60850320
60850305
60850315

Table S15. Primer sequences used to generate CRSs.

| Enhancer | Forward Primer | Reverse Primer |
|---|---|---|
| *BCL2_1* | TCCTCACACCACTTATCCTCCTTGCCTGTCCCTGGCG | GTCTGGGGTGGGGTGGG |
| *BCL2_2* | TCCAAACAGCTGCAGTTCA | GTGAGGAACCAAGAGAGG |
| *BCL2_3* | ACCAACATTCCCACAATGCCCAGTGGAAA | AGAAGGGTGGTGAGAGAGGTGTGTGGTTATTCTTGCT |
| *ADGRG6*_rs1040525 | CATGTCTATAAAATAAAGAG GGAAATTAATTTGACTATTCAGT | TGGATAATTAGGAAGAGCAATGATAAAATAAAGCAAAA |
| *BNC2*_rs10738446 | AATACCCTCGCCTGTATT | AGAAATAAGTTAGTTCTTGATCCTAAGGAGAGAAAAA |
| *FTO*_rs7201850 | GACACTGAACTATTCA | ATCTTCAACACCAAT |
